# Nucleus reuniens is required for encoding and retrieving precise, hippocampal-dependent contextual fear memories in rats

**DOI:** 10.1101/340190

**Authors:** Karthik R. Ramanathan, Reed L. Ressler, Jingji Jin, Stephen Maren

**Author notes:** Corresponding author: Stephen Maren, Department of Psychological and Brain Sciences, TAMU 4235, Texas A&M University, College Station, TX 77843-3474. Author contribution:* KR and SM designed the experiments, analyzed the data and wrote the manuscript. KR, JJ and RR collected the data. Data availability:* The datasets generated during and/or analyzed during the current study are available from the corresponding author upon request.

## Abstract

The nucleus reuniens (RE) is a ventral midline thalamic nucleus that interconnects the medial prefrontal cortex (mPFC) and hippocampus (HPC). Considerable data indicate that HPC-mPFC circuits are involved in contextual and spatial memory; however, it is not clear whether the RE mediates the acquisition or retrieval of these memories. To examine this question, we inactivated the RE with muscimol before either the acquisition or retrieval of Pavlovian fear conditioning in rats; freezing served as the index of fear. We found that RE inactivation before conditioning impaired the acquisition of contextual freezing, whereas inactivation of the RE prior to retrieval testing increased the generalization of freezing to a novel context; inactivation of the RE did not affect either the acquisition or expression of auditory fear conditioning. Interestingly, contextual conditioning impairments were absent when retrieval testing was also conducted after RE inactivation. Contextual memories acquired under RE inactivation were hippocampal-independent, insofar as contextual freezing in rats conditioned under RE inactivation was insensitive to intra-hippocampal infusions of the NMDA receptor antagonist, D,L-amino-5-phosophonovaleric acid (APV). Together, these data reveal that the RE supports hippocampal-dependent encoding of precise contextual memories that allow discrimination of dangerous from safe contexts. When the RE is inactive, however, alternate neural systems acquire an impoverished contextual memory that is only expressed when the RE is offline.

**SIGNIFICANCE STATEMENT:** The midline thalamic nucleus reuniens (RE) coordinates communication between the hippocampus and medial prefrontal cortex, brain areas critical for contextual and spatial memory. Here we show that temporary pharmacological inactivation of RE impairs the acquisition and precision of contextual fear memories after Pavlovian fear conditioning in rats. However, inactivating the RE prior to retrieval testing restored contextual memory in rats conditioned after RE inactivation. Critically, we show that imprecise contextual memories acquired under RE inactivation are learned independently of the hippocampus. These data reveal that the RE is required for hippocampal-dependent encoding of precise contextual memories to support the discrimination of safe and dangerous contexts.

## Introduction

The nucleus reuniens (RE) is a ventral midline thalamic nucleus that interconnects the medial prefrontal cortex (mPFC) and hippocampus (HPC) and plays a critical role in both memory and emotion (Jin and Maren, 2015a; McKenna and Vertes, 2004; Preston and Eichenbaum, 2013; Vertes et al., 2006). Recent work has shown that lesions or inactivation of the RE reduces synchrony between the mPFC and HPC and produces deficits in tasks that require coordinated activity between the HPC and mPFC (Davoodi et al., 2011; Hembrook and Mair, 2011; Hembrook et al., 2012; Cholvin et al., 2013;). For example, RE inactivation impairs performance in spatial working memory task that requires delayed alternation in a T-maze, and these memory deficits are accompanied by reductions in theta coherence between the mPFC and HPC (Layfield et al., 2015; Hallock et al., 2016).

Deficits in spatial working memory associated with RE inactivation may be due to deficits in contextual processing associated with impaired HPC-mPFC interaction (Cassel and Pereira de Vasconcelos, 2015). Consistent with this possibility, chronic inhibition of synaptic transmission in the RE made with a virally expressed tetanus toxin (TetTox) disrupts contextual processing during Pavlovian fear conditioning in mice (Xu and Südhof, 2013). Specifically, permanent RE inactivation increased the generalization of fear to a novel context after contextual fear conditioning. These results suggest that the RE is involved in encoding or consolidating (Vetere et al., 2017; Troyner et al., 2018) precise context representations that normally limit the generalization of fear from the conditioning context to other, dissimilar contexts.

Once learned, however, the retrieval of contextual memories does not appear to require the RE (Xu and Südhof, 2013). Yet there is considerable data in both animals and humans indicating that HPC-mPFC interactions are involved in the retrieval of spatial and contextual memories (Orsini et al., 2011; Preston and Eichenbaum, 2013; Schlichting and Preston, 2016; Wang et al., 2016; Marek et al., 2018). Indeed, recent work indicates that reversible optogenetic inhibition of the hippocampus yields deficits in both the encoding and retrieval of contextual fear memories (Goshen et al., 2011; Bernier et al., 2017). Moreover, RE inactivation produces robust impairments in spatial working memory (Griffin, 2015; Layfield et al., 2015; Hallock et al., 2016). Hence, intact acquisition and expression of contextual freezing in rats with permanent lesions of the RE (Xu and Südhof, 2013) may be due to compensation by alternate neural systems, which has been observed after hippocampal lesions, for example (Maren et al., 1997). Insofar as the RE plays a role in supporting HPC-mPFC interactions, we hypothesize that reversible inactivation of the RE will impair both the acquisition and expression of contextual fear.

To examine this question, we temporarily inactivated RE using intra-cranial infusions of muscimol (MUS, a GABA_A_ agonist) during either the acquisition or retrieval (or both) of Pavlovian fear conditioning in rats. Retention tests were conducted in both the conditioning context and an alternate context to assess the influence of RE inactivation on context discrimination and generalization; both contextual and auditory freezing were assessed. Importantly, we included control groups in our design to determine whether state-dependent generalization deficits account for performance impairments in MUS-treated rats. Consistent with our hypothesis, we found that the RE is involved in both the encoding and retrieval of context representations that support contextual discrimination. Interestingly, deficits in contextual freezing after fear conditioning under RE inactivation could be rescued by inactivating the RE during retrieval testing. However, the contextual memories acquired under RE inactivation were hippocampal-independent, insofar as contextual freezing in rats conditioned under RE inactivation was insensitive to intra-hippocampal infusions of D,L-amino-5-phosophonovaleric acid (APV). This supports the view that the RE is a component of a hippocampal memory system that prioritizes the encoding of a configural representation of context that ultimately overshadows its underlying elements.

## Materials and Methods

### Subjects

One hundred and seventy experimentally naïve adult male Long-Evans rats (Blue-Spruce; 200– 224 g; 50–57 days old; RRID:RGD_5508398) were obtained from a commercial supplier (Envigo, Indianapolis, IN). The rats were individually housed in cages within a temperature- and humidity-controlled vivarium and kept on a 14:10 h light:dark cycle (lights on at 07:00 hours) with *ad libitum* access to food and water. All experiments took place during the light phase of the cycle. Rats were handled for one minute per day for 5 days to habituate them to the experimenter before they underwent surgery. All experiments were conducted at Texas A&M University with approval from its Animal Care and Use Committee.

### Surgical procedures

One week before behavioral testing, rats were anesthetized with isoflurane (5% for induction, ~2% for maintenance) and placed into a stereotaxic instrument (Kopf Instruments). An incision was made in the scalp and the head was leveled by placing bregma and lambda in the same horizontal plane. Small holes were drilled in the skull to affix three jeweler’s screws and to allow implantation of guide cannulas (coordinates relative to bregma) targeting either RE or dorsal hippocampus (DH). For Experiment 1, RE was targeted using a single guide cannula (8 mm, 26 gauge; Plastics One) implanted at a 10° angle on the midline (A/P: -2.0 mm, M/L: -1.0 mm, D/V: -6.7 mm from dura). For Experiment 2, three guide cannulas were implanted to target RE (single midline) and DH (bilateral) in the same animal. The DH was targeted using two guide cannulas (4 mm, 26 gauge; Plastics One) implanted at a 20° angle (A/P: -3.7 mm, M/L: ±3.5 mm, D/V: -3.2 mm from dura). The guide cannulas were affixed to the skull with dental cement and stainless-steel dummy cannulas (31 gauge, 9 mm for the RE and 5 mm for the DH; Plastics One) were inserted into the guides. Rats were allowed to recover for 7 d after surgery before behavioral testing; dummy cannulae were replaced twice during this period.

### Drug infusions

For intracranial microinfusions, rats were transported to the laboratory in 5-gallon white buckets filled with a layer of bedding. The dummies were removed and stainless-steel injectors (31 gauge, 9 mm for RE and 5 mm for DH) connected to polyethylene (PE) tubing were inserted into the guide cannulas for intracranial infusions. The PE tubing on each injector was connected to a Hamilton syringe (10 µl), which was mounted on an infusion pump (Kd Scientific). Infusions were monitored by the movement of an air bubble that separated the drug or saline solutions from distilled water within the PE tubing. All infusions were made approximately 10 min before either the conditioning or retrieval test sessions. Muscimol (MUS; 0.1 µg/µl) and APV (10 µg/µl) were dissolved in sterile saline (SAL) and infused at a rate of 0.1 µl/min for 3 min (0.3 µl total; 0.03 µg MUS in RE and 3 µg APV per hemisphere in DH); the injectors were left in place for 2-3 min for diffusion. After the infusions, clean dummies were inserted into the guide cannulae and the animals were transported to the conditioning chambers for the behavioral sessions.

### Behavioral apparatus

Sixteen identical rodent conditioning chambers (30 × 24 × 21 cm; Med-Associates, St Albans, VT) housed in sound-attenuating cabinets were used in all behavioral sessions. Each chamber consisted of two aluminum sidewalls, a Plexiglas ceiling and rear wall, and a hinged Plexiglas door. The floor consisted of 19 stainless steel rods that were wired to a shock source and a solid-state grid scrambler (Med-Associates) for the delivery of footshock. A speaker mounted outside the grating of one aluminum wall was used to deliver auditory stimuli. Additionally, ventilation fans in the cabinets and a house light in each chamber were used to create distinct contexts. Each conditioning chamber rested on a load-cell platform that was used to record chamber displacement in response to each rat’s motor activity. Load-cell voltages were digitized and recorded on a computer using Threshold Activity software (Med-Associates). For each chamber, load-cell voltages are digitized at 5 Hz, yielding one observation every 200 ms. Freezing was quantified by computing the number of observations for each rat that had a value less than the freezing threshold (load-cell activity = 10). Freezing was only scored if the rat is immobile for at least 1 sec. Environmental stimuli were manipulated within two different laboratory rooms housing the conditioning chambers (8 chambers/room) to generate two distinct contexts. For context A, a 15-W house light within each chamber was turned on and the overhead fluorescent room lights remained on. Ventilation fans (65 dB) were turned on, cabinet doors were left open, and the chambers were cleaned with 1% ammonium hydroxide. Rats were transported to context A in white plastic boxes. For context B, the house lights were turned off and a overhead red fluorescent room light was turned on. The cabinet doors were closed and the chambers were cleaned with 1.5% acetic acid. Rats were transported to context B in black plastic boxes.

### Behavioral procedures: Experiment 1

The experiment was run in two replications with half of the animals undergoing a contextual conditioning fear procedure (*n* = 64) and the other half undergoing an auditory fear conditioning procedure (*n* = 64). There were no statistical differences (all *F*’s < 1.2; *p* > 0.3) across these replications in the effects of RE inactivation on the acquisition or expression of freezing during conditioning or the context retrieval tests, so the data from these sessions were collapsed. Conditioned freezing to the tone CS was assessed in only the cohort that underwent auditory fear conditioning.

Approximately one week after surgery, the animals were randomly assigned to one of four groups: SAL-SAL, SAL-MUS, MUS-SAL and MUS-MUS. On day 1 rats received microinfusions of either MUS or SAL and were subjected to either contextual or auditory fear conditioning in context A. The conditioning session consisted of a 3-min baseline followed by 5 footshock unconditioned stimuli (US; 1 mA, 2 sec) separated by 70-s inter-trial intervals (ITI). In half of the animals, the US was preceded by a 10-sec auditory conditioned stimulus (CS; 2 kHz, 80 dB). Twenty-four hours later, rats again received microinfusions of MUS or SAL and were placed in either the conditioning context (A) or a novel context (B) for a 10-min stimulus-free retrieval test to assess conditioned freezing and its generalization, respectively. Each rat was tested in each context in a counterbalanced fashion across two days and the infusion group assignment was the same for each test. Rats that received auditory fear conditioning were tested identically, except that five CS-alone test trials (30-sec ISI) were delivered after the 10-min baseline (generalization test) period in context B.

### Behavioral procedures: Experiment 2

This experiment was run in two replications. After recovery from surgery, animals were assigned to one of four groups: DH_SAL_-RE_SAL_, DH_SAL_-RE_MUS_, DH_APV_-RE_SAL_ and DH_APV_-RE_MUS_. Animals received either SAL or APV infusions into the DH and either SAL or MUS into the RE prior to conditioning; all animals were then tested after an infusion of SAL or MUS into the RE (the RE drug infusion during testing was the same as that during conditioning). This design permitted us to determine whether contextual conditioning under RE inactivation requires NMDA receptors in the DH. After DH and RE infusions on day 1, the conditioning session consisted of a 3-min baseline followed by 5 footshocks unconditioned stimuli (US; 1 mA, 2 sec) separated by 70-s inter-trial intervals (ITI). Forty-eight hours after conditioning, rats again received microinfusions of MUS or SAL in RE and were placed in the conditioning context (A) for a 10-min stimulus-free retrieval test to assess contextual freezing.

### Histological procedures

Upon completion of the experiment, rats were overdosed with sodium pentobarbital (Fatal Plus, 100 mg/kg) and perfused transcardially with 0.9% saline followed by 10% formalin. The brains were extracted from the skull and post-fixed in a 10% formalin solution for 24 h followed by a 30% sucrose-formalin solution where they remained for a minimum of 72h. After the brains were fixed, coronal sections (40 µm thickness) were made on a cryostat (−20 °C), mounted on subbed microscope slides, and stained with thionin (0.25%) to visualize cannula placements (see Figure 1A for representative cannula placement).

**Figure 1.**
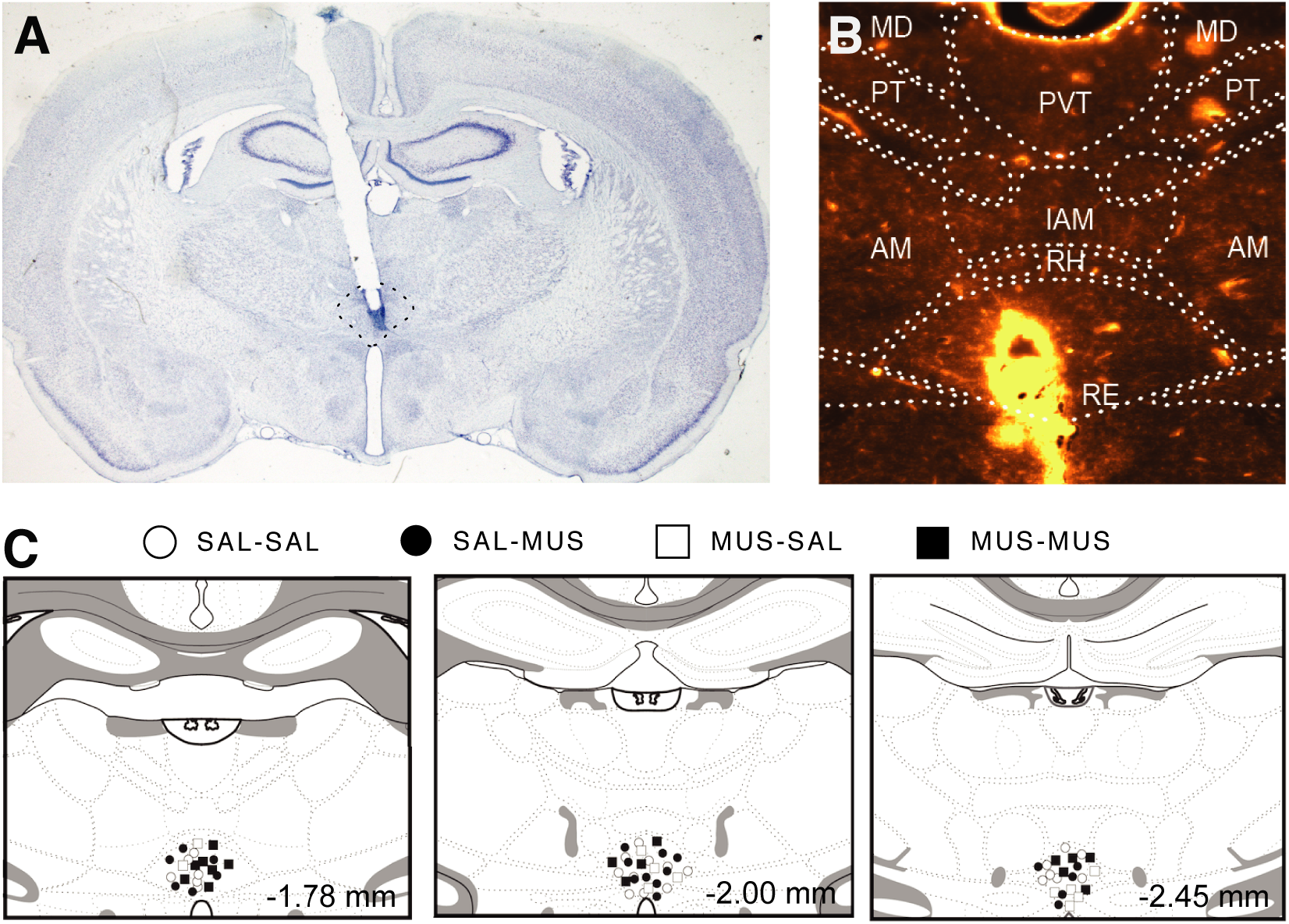
Cannula placements in nucleus reuniens (RE). *A*, Representative thionin-stained coronal section showing a midline cannula placement in the RE. *B*, Representative darkfield image showing diffusion of fluorescent muscimol (0.3 µl; TMR-X) in the RE. *C*, Cannula placements for all subjects that were included in the analysis across three different levels along the anterior-posterior axis. The distribution of cannula placements within the RE was similar across all of the groups.

### Experimental design and statistical analysis

In Experiment 1, rats (*n* = 128) were randomly assigned to a 2 X 2 factorial design with variables of drug condition (SAL or MUS infusions into RE) prior to fear conditioning and/or retrieval testing. This design resulted in four experimental groups: SAL-SAL, SAL-MUS, MUS-SAL and MUS-MUS. Half of the animals underwent contextual fear conditioning (*n* = 64; *n* = 16/group) and the other half underwent auditory fear conditioning (*n* = 64; *n* = 16/group). Ultimately, the contextual freezing data were collapsed across these experiments resulting 32 subject per group. During the course of the experiment, three rats died or were sacrificed before completion of the experiment and were excluded from the analysis. Of the remaining animals, fourteen were excluded due to poor cannula placements resulting in the following final group sizes: SAL-SAL (*n* = 29), SAL-MUS (*n* = 27), MUS-SAL (*n* = 31), MUS-MUS (*n* = 24) for the conditioning session and context retrieval tests. Freezing to the tone was assessed in roughly half the number of subjects: SAL-SAL (*n* = 16), SAL-MUS (*n* = 13), MUS-SAL (*n* = 16), MUS-MUS (*n* = 11).

In Experiment 2, rats (*n* = 42) were assigned to a 2 X 2 factorial design with variables of DH drug condition prior to fear conditioning (SAL or APV) and RE drug condition prior to conditioning and retrieval testing (SAL-SAL or MUS-MUS). This design resulted in four experimental groups: DH_SAL_-RE_SAL_ (*n* = 9), DH_APV_-RE_SAL_ (*n* = 13), DH_SAL_-RE_MUS_ (*n* = 8), DH_APV_ - RE_MUS_ (*n* = 12). Two rats died or were sacrificed before completion of the experiment; of the remaining 40 animals, eight were excluded due to poor cannula placements or inadvertent lesions associated with cannula placement or drug infusions. One animal that exhibited average freezing that was greater than two standard deviations above its group mean was excluded resulting in the following final group sizes: DH_SAL_-RE_SAL_ (*n* = 9), DH_APV_- RE_SAL_ (*n* = 9), DH_SAL_-RE_MUS_ (*n* = 7), DH_APV_ - RE_MUS_ (*n* = 7).

All behavioral data (mean ± SEM) are represented by the average percentage of freezing behavior during one-minute intervals during the conditioning session and retrieval tests. For those animals receiving a tone retrieval test, freezing was averaged across both the auditory CS and the subsequent 30-sec ISI (CS+ITI). Freezing during the CS+ITI period is highly correlated with freezing to the CS itself and is less susceptible to competition by active CS-elicited orienting responses. For the conditioning and retrieval test sessions, the behavioral data were analyzed with a three-way mixed-model analysis of variance (ANOVA) with between-subject factors of drug condition during conditioning and testing and within-subject factors test minute. Analyses of contextual discrimination performance consisted of a two-way mixed-model ANOVA with a between-subject factor of drug condition during conditioning and testing and a within-subject factor of test context [conditioning (A) or novel (B)]. Lastly, discrimination ratio and auditory freezing data in Experiment 1 and contextual test performance in Experiment 2 were the freezing data were averaged across the test minutes and these data were analyzed with two-way ANOVAS with between-subject factors of drug condition during conditioning and retrieval testing. Post-hoc comparisons in the form of Fisher’s protected least significant difference (PLSD) tests were performed after a significant overall *F*-ratio in the ANOVA (*α* = 0.05 for both main effects and interactions). A sample size of eight rats per group is sufficient to detect the effect sizes anticipated in the present experiments (G*Power), and the final group sizes in each experiment resulted from two independent replications. Statistical analyses were performed with StatView 5.0.1 (SAS Institute, Inc) running under an open-source PowerPC Apple MacOS emulator (SheepShaver; https://sheepshaver.cebix.net).

## Results

### Inactivation of RE impairs acquisition of contextual conditioning and generalization of conditioned freezing

Figure 1 (A and B) illustrates the spread of fluorescently labeled muscimol (TMR-X) in the RE along with an illustration of cannula placements in all the animals included in the analyses (Figure 1C). During the fear conditioning session, rats receiving SAL or MUS infusions into the RE exhibited low levels of freezing behavior prior to the first conditioning trial and increased their freezing behavior across the conditioning trials (Figure 2A); there were no differences between MUS- and SAL-treated in the levels of conditioned freezing. These observations were confirmed in a one-way repeated measures ANOVA that revealed a significant main effect of training trial [*F*(5, 545) = 166.9; *p* < 0.0001] with neither a main effect of drug [*F*(1,109) = 3.56; *p* = 0.06] nor a drug x trial interaction [*F*(5,545) = 0.78; *p* = 0.57]. Hence, RE inactivation did not affect the expression of post-shock freezing during the acquisition of Pavlovian fear conditioning.

**Figure 2.**
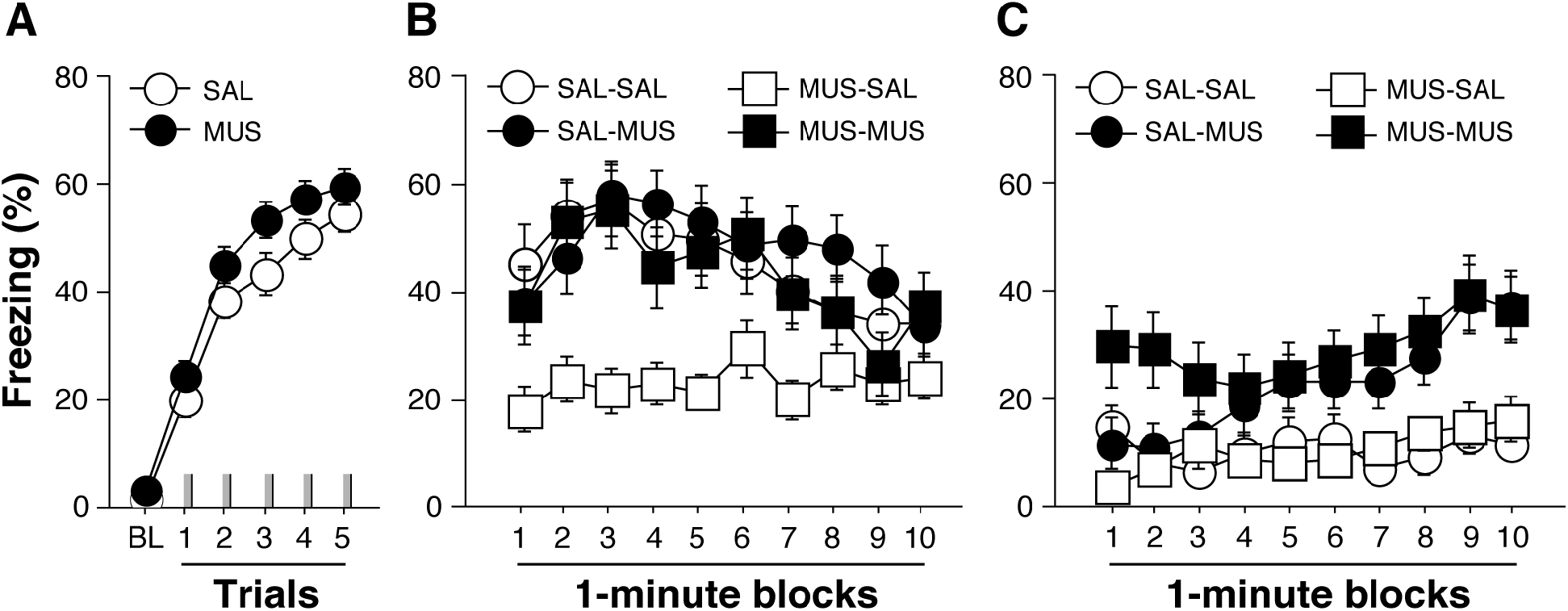
Muscimol inactivation of the RE results in a state-dependent impairment in contextual fear conditioning. *A*, Percentage of freezing during the 3-min baseline (BL) and 1-min ITI (intertrial interval) after each conditioning trial (indicated by gray hatch marks on the X-axis) in animals infused with either saline (SAL) or muscimol (MUS) in the RE. *B*, Percentage of freezing during the 10-minute retrieval test in the conditioning context. Animals conditioned after SAL or MUS infusions received the retrieval test after either SAL or MUS infusions in a factorial design that yielded four groups SAL-SAL, MUS-SAL, SAL-MUS, MUS-MUS. *C*, Percentage of freezing during the 10-minute retrieval test in a novel context (the order of the retrieval tests in *B* and *C* was counterbalanced); the groups are the same as those described in *B*. All data are shown as means ± SEMs.

Twenty-four and forty-eight hours later, all animals were again infused with either SAL or MUS and placed in either the conditioning context (A) or a novel context (B) for a 10-min retrieval test; the test order in the two contexts was counterbalanced and the infusion assignment for each test was the same. Importantly, this design allowed us to assess the state-dependent effects of RE inactivation on the acquisition, expression, and generalization of conditioned freezing. As shown in Figure 2B, rats conditioned after MUS infusions into the RE (MUS-SAL group) exhibited a substantial impairment in conditioned freezing in the conditioning context. Interestingly, this impairment was absent in animals both conditioned and tested after MUS infusions into the RE (MUS-MUS group). The relatively high level of freezing behavior in the MUS-MUS rats was not simply due to a nonspecific increase in freezing caused by RE inactivation insofar as SAL-MUS animals were no different from SAL-SAL or MUS-MUS animals. All of these observations were confirmed in a two-way repeated measures ANOVA that revealed significant main effects for conditioning drug [*F*(1, 107) = 8.28; *p* = 0.004], testing drug [*F*(1, 107) = 6.04; *p* = 0.02], and time [*F*(1, 107) = 8.90; *p* < 0.0001]. Moreover, there was a trend towards a significant interaction between conditioning and testing drug conditions [*F*(1, 107) = 3.52; *p* = 0.06] (Figure 2B). Post-hoc comparisons revealed that freezing among rats in the MUS-SAL group was significantly lower than that in the SAL-SAL (*p* = 0.0007), SAL-MUS (*p* = 0.0002) and MUS-MUS (*p* = 0.003) groups, which did not differ from one another. This reveals that RE inactivation impairs the acquisition of contextual fear conditioning and does so in a state-dependent manner; testing animals in the same state under which they were conditioned resulted in high levels of conditioned freezing.

In contrast, when the animals were tested for their generalization of fear to a novel context a different pattern of results emerged. As shown in Figure 2C, rats conditioned after MUS inactivation of the RE exhibited similar and low levels of freezing relative to animals conditioned under SAL. However, MUS infusions prior to the generalization test increased conditioned freezing. Interestingly, animals conditioned and tested under MUS (MUS-MUS) exhibited the highest level of freezing, and this was manifest early in the test session. Rats conditioned under SAL and tested under MUS (SAL-MUS) showed low levels of freezing at the beginning of the test that increased over the course of the test to approach those in the MUS-MUS group. These observations were confirmed in a two-way repeated measures ANOVA that revealed a significant main effect of test drug [*F*(1,107) = 22.78; *p* < 0.0001] and time [*F*(9,963) = 7.65; *p* < 0.0001]; there was no interaction between conditioning and test drug [*F*(1,107) =1.1; *p* = 0.29]. However, there was a significant interaction between test drug and time [*F*(9,963) = 2.31; *p* = 0.014], which indicates that MUS increased the generalization of fear (SAL-MUS), and served as a retrieval cue to support contextual freezing in rats conditioned after MUS infusions in the RE (MUS-MUS). Importantly, increases in freezing were not due to nonspecific effects of MUS on locomotion, insofar as MUS did not decrease locomotor (i.e., load-cell) activity during the pre-trial baseline period on the conditioning day, nor did it affect shock-elicited activity (data not shown).

### RE inactivation impairs contextual discrimination

Because animals received the same drug infusions across both of the context tests, we were able to assess contextual discrimination using a within-subjects analysis. As shown in Figure 3A, rats in all the groups exhibited a contextual discrimination and exhibited lower levels of freezing in context B compared to context A. Consistent with this observation, a three-way repeated measures ANOVA with between-subject variables of conditioning and testing drug condition and a within-subject variable of test context revealed a significant main effect of test context [*F*(1,107) = 80.48; *p* < 0.0001]. Furthermore, there was a significant main effect of test drug [*F*(1,107) = 17.42; *p* < 0.0001], but not conditioning drug [*F*(1,107) = 2.3; *p* = 0.13], although there was a trend towards a significant interaction between the drug conditions [*F*(1,107) = 3.48; *p* = 0.06]. Close examination of these data reveals that the SAL-SAL animals exhibited the highest level of contextual discrimination relative to all the other groups. To examine this possibility more closely, we calculated a discrimination index by computing the ratio between the differences in the average freezing across the 10-min tests (context A - context B) divided by the total freezing in each context (context A + context B). As shown in Figure 3B, MUS infusions into RE resulted in lower discrimination scores whether the infusions occurred before conditioning or retrieval testing. A factorial ANOVA revealed a trend towards a significant main effect of conditioning drug [*F*(1,107) = 3.62; *p* = 0.06] and a significant main effect of drug during testing [*F*(1,107) = 6.83; *p* = 0.01]; there was no significant interaction between these factors [*F*(1,107) = 0.16; *p* = 0.69]. Hence, it appears that RE inactivation reduces contextual discrimination when infused either before conditioning or retrieval testing.

**Figure 3.**
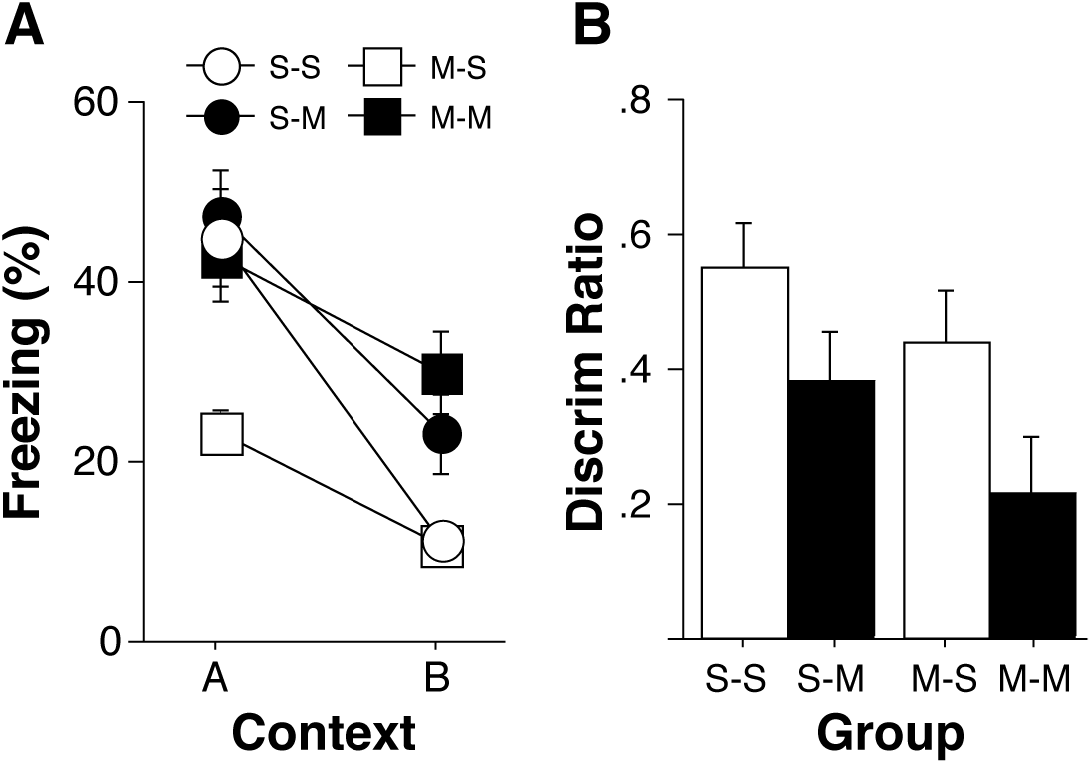
Muscimol inactivation of the RE impairs contextual discrimination. *A*, Average percentage of freezing across the 10-minute retrieval tests in the conditioning context (context A) and the novel context (context B) shown in Figure 2. The order of the two retrieval tests was counterbalanced and animals trained under saline (S) or muscimol (M) were tested in both contexts A and B under after either saline or muscimol infusions to yield the four groups described in Figure 2 (e.g., S-S, S-M, M-S, and M-M). *B*, Average discrimination ratios [(context A - context B)/(context A + context B)] calculated on freezing behavior across the 10-minute tests shown in *A*. All data are shown as means ± SEMs.

### RE inactivation does not impair acquisition or expression of freezing to an auditory CS

Of the subset of animals that received auditory fear conditioning, we determined whether inactivation of RE causes deficits in acquisition or expression of fear to the tone CS. As shown in Figure 4A and B, conditional freezing to the tone was similar in all of the groups. Two-way repeated measures ANOVA revealed no significant main effects or interactions of drug on conditioning [*F*(1,52) = 2.6; *p* = 0.12] or testing [*F* (1,52) = 1.77; *p* = 0.19] across the entire test. Moreover, average freezing during the CS trials (Figure 4B) did not differ between the groups [conditioning, *F*(1,52) = 3.47; *p* = 0.07; testing, *F* (1,52) = 1.05; *p* = 0.31]. Overall, these results indicate that inactivation of RE does not cause deficits in either the acquisition or retrieval of fear to an auditory CS.

**Figure 4.**
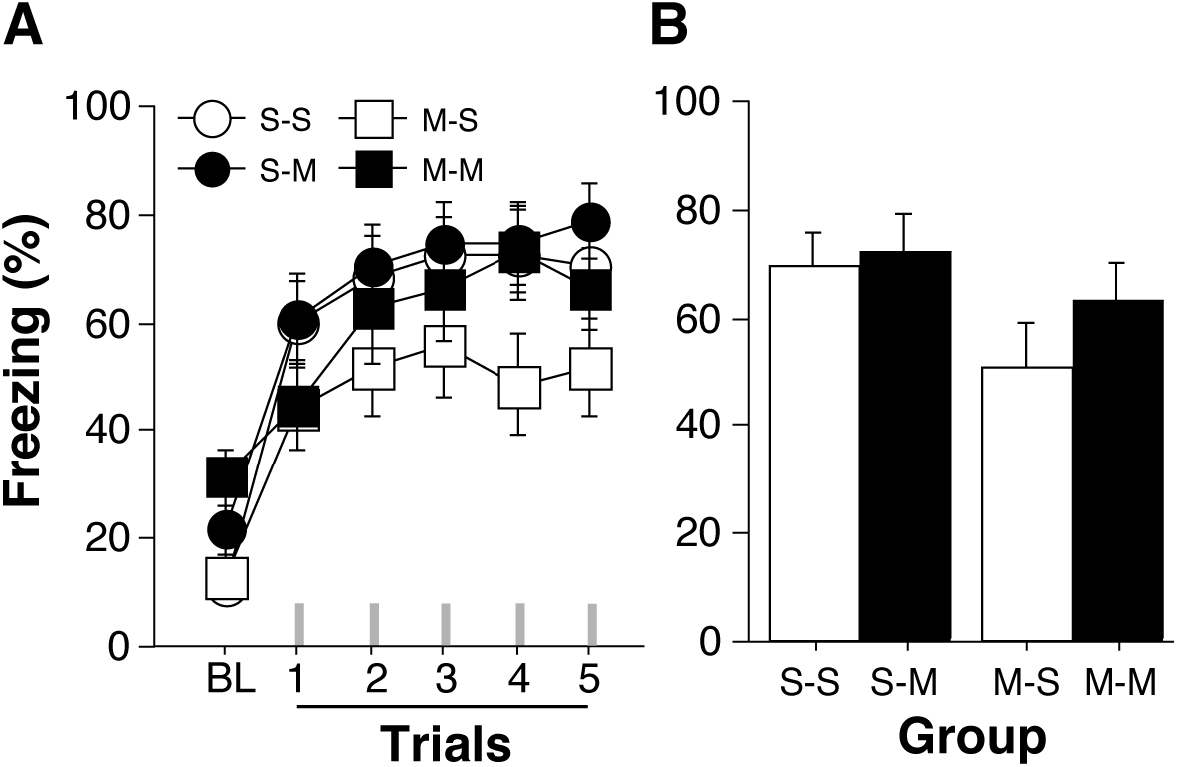
Muscimol inactivation of RE did not impair the acquisition or expression of freezing to an auditory conditioned stimulus (CS). *A*, Average percentage of freezing during a 10-min stimulus free baseline (BL) and after five tone presentations (indicated by the grey hatch marks on the x-axis) during an auditory retrieval test. Freezing during each trial represents the average freezing during the CS and intertrial interval. Animals received either saline (S) or muscimol (M) infusions prior to conditioning and retrieval testing in a factorial design to yield four groups (S-S, M-S, S-M, M-M). *B*, Average percentage of freezing across all five CS trials shown in *A*. All data are shown as means ± SEMs.

### Contextual memories acquired under RE inactivation are insensitive to hippocampal NMDA receptor antagonism

Animals conditioned and tested under RE inactivation exhibited high levels of freezing in the conditioning context, suggesting that they had acquired a contextual memory. Because considerable evidence indicates that RE inactivation impairs hippocampal-dependent memory encoding (Layfield et al., 2015; Hallock et al., 2016), contextual learning in rats conditioned under RE inactivation may not require hippocampal synaptic plasticity (Stiedl et al., 2000; Quinn et al., 2005; Czerniawski et al., 2012). To test this possibility, we examined whether intra-hippocampal infusions of the NMDA receptor antagonist, APV, impair contextual conditioning when training occurs under simultaneous RE inactivation.

Representative cannula placements in RE or DH along with an illustration of cannula placements in all the animals included in the analyses are shown in Figure 5. During fear conditioning, all rats exhibited low levels of freezing behavior prior to the first conditioning trial and increased their freezing behavior across the conditioning trials (Figure 6A). There were no differences between the groups in the levels of conditioned freezing. A one-way repeated measures ANOVA revealed a significant main effect of conditioning trial [*F*(5,140) = 32.79; *p* < 0.0001] with neither a main effect of drug in RE [*F*(1,28) = 0.005; *p* = 0.94] and drug in DH [*F*(1,28) = 1.81; *p* = 0.19] nor a drug x trial interaction [*F*(1,28) = 1.17; *p* = 0.29]. Hence, neither MUS infusions into RE nor APV into DH affected post-shock freezing during the acquisition of Pavlovian fear conditioning.

**Figure 5.**
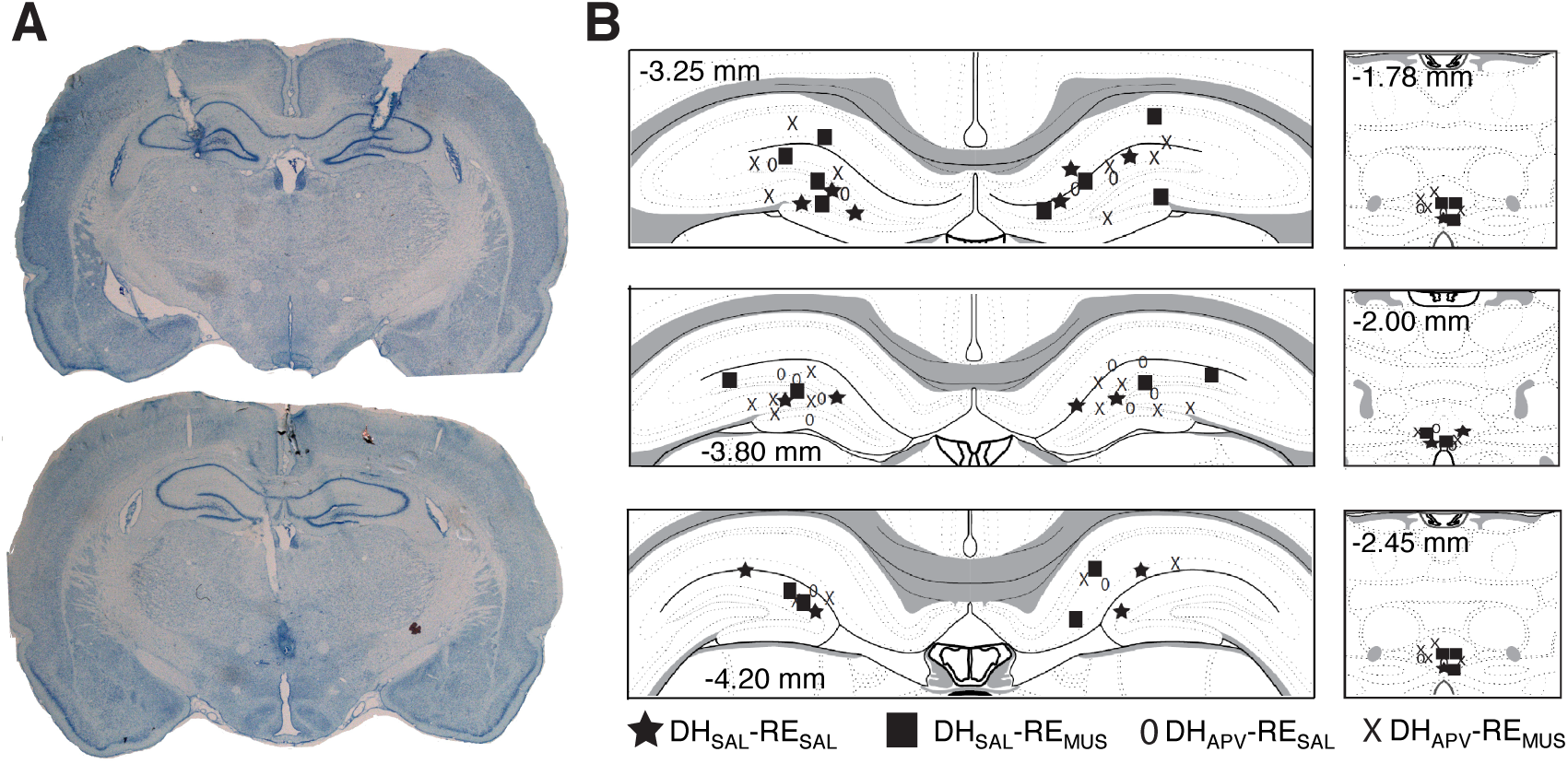
Cannula placements in nucleus reuniens (RE) and dorsal hippocampus (DH). *A*, Representative thionin-stained coronal sections showing a midline cannula placement in the RE (left) and DH (right). *B*, Cannula placements for all subjects that were included in the analysis across three different levels along the anterior-posterior axis in both RE (left) and DH (right). The distribution of cannula placements within the RE and DH was similar across all of the groups.

**Figure 6.**
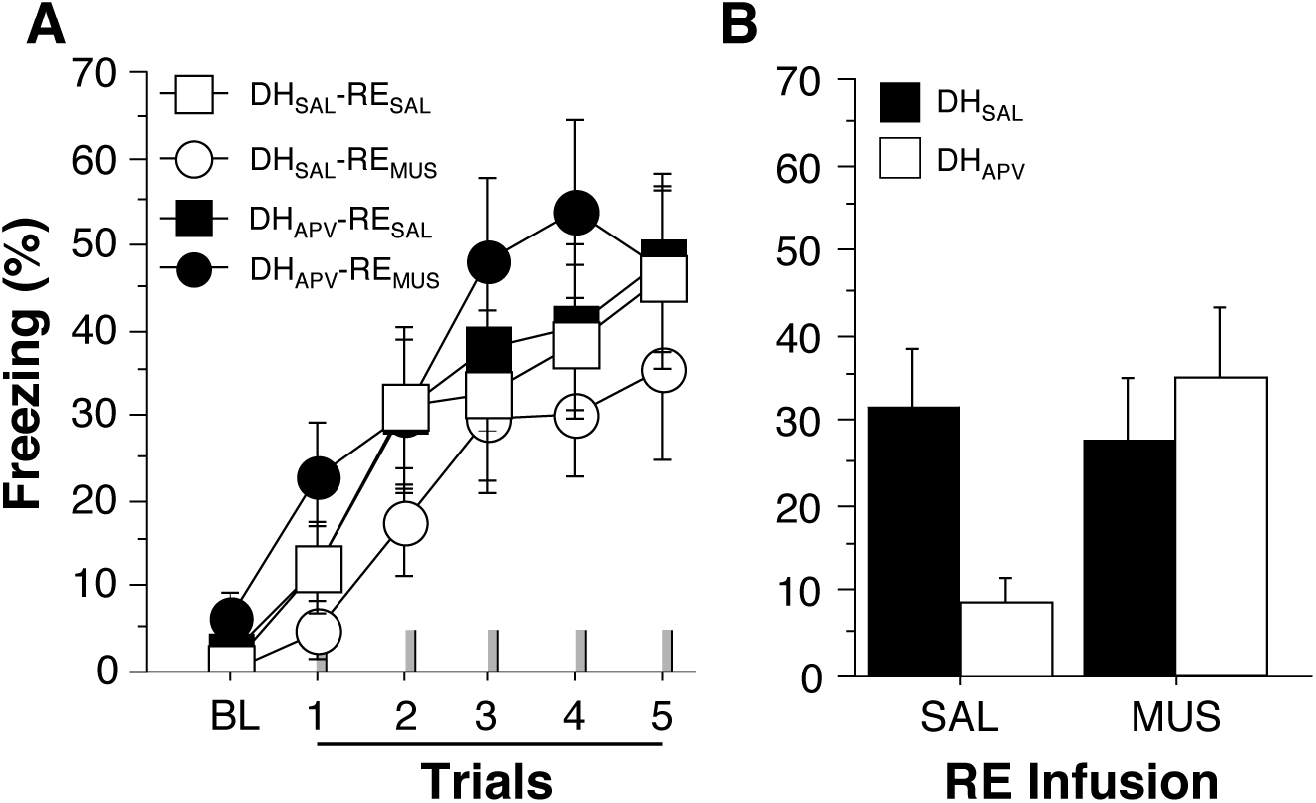
Contextual conditioning under RE inactivation is insensitive to NMDA receptor antagonism in the DH. *A*, Percentage of freezing during the 3-min baseline (BL) and 1-min ITI (intertrial interval) after each conditioning trial (indicated by gray hatch marks on the X-axis) in animals infused with either saline (SAL) or muscimol (MUS) in the RE and APV or saline (SAL) infused in DH. *B*, Percentage of freezing during the 10-minute retrieval test in the conditioning context. Animals conditioned after SAL or MUS infusions in RE received the same drug infusions in RE prior to the retrieval test in a factorial design that yielded four DH_SAL_-RE_SAL_, DH_SAL_-RE_MUS_, DH_APV_-RE_SAL_ and DH_APV_-RE_MUS_. SAL or APV indicates drug infusions in DH prior to fear conditioning and SAL or MUS indicate the RE infusions prior to both conditioning and the retrieval test. All data are shown as means ± SEMs

Forty-eight hours later, all animals were again infused with either SAL or MUS in RE and placed in the conditioning context (A) for a 10-min retrieval test (the drug infused prior to retrieval testing was identical to that infused prior to conditioning); there were no DH infusions prior to retrieval test. As shown in Figure 6B, rats conditioned after APV infusions into the DH and SAL infusions into the RE (DH_APV_-RE_SAL_) exhibited a substantial impairment in conditioned freezing in the conditioning context relative to animals receiving SAL infusions into both DH and RE (DH_SAL_-RE_SAL_). Interestingly, this impairment was absent in animals that received MUS infusions to RE (DH_APV_-RE_MUS_) who froze at levels similar to that in controls (DH_SAL_-RE_SAL_ and DH_SAL_RE_MUS_). The relatively high level of freezing behavior in the DH_APV_-RE_MUS_ animals was not simply due to a nonspecific increase in freezing caused by drug infusions in RE insofar as the level of freezing in DH_APV_-RE_MUS_ group was similar to that in animals conditioned under RE inactivation alone (DH_SAL_-RE_MUS_) and saline controls (DH_SAL_-RE_SAL_). These observations were confirmed in a two-way repeated measures ANOVA that revealed was a significant interaction between RE and DH drug conditions [*F*(1,28) = 5.91; *p* = 0.02]; there was no main effect of RE drug condition [*F*(1,28) = 3.11; *p* = 0.09] or DH drug condition [*F*(1,28) = 1.52; *p* = 0.23]. Post-hoc comparisons revealed that freezing among rats in the DH_APV_-RE_SAL_ group was significantly lower than that in the DH_SAL_-RE_SAL_ (*p* = 0.0098), DH_SAL_-RE_MUS_ (*p* = 0.0432) and DH_APV_-RE_MUS_ (*p* = 0.0061) groups, which did not differ from one another. This reveals that whereas contextual conditioning normally requires hippocampal NMDA receptors, contextual memories acquired after inactivation of the RE do not require DH NMDA receptors.

## Discussion

The present study examined the role of the RE in the acquisition, expression, and generalization of conditioned freezing in rats. Inactivation of RE prior to fear conditioning impaired contextual freezing in the conditioning context. Importantly, this deficit could be completely rescued by testing animals under RE inactivation, indicating that the contextual memory acquired under RE inactivation can be expressed, but only when the RE is offline. Memories acquired or expressed under RE inactivation were less precise, insofar as RE inactivation increased generalization of contextual fear to a novel context. Lastly, contextual memories acquired under RE inactivation were not impaired by NMDA receptor antagonism in the DH, revealing that non-hippocampal systems acquire contextual memory in the absence of the RE. Collectively, these data reveal that the RE is required for hippocampal-dependent encoding of precise contextual memories to support the discrimination of safe and dangerous contexts.

The discovery that RE inactivation impairs the acquisition of precise context memories is consistent with prior work showing that TetTox inhibition of synaptic inputs to the RE, particularly from the PFC, leads to increased generalization of contextual fear without affecting fear to an auditory CS (Xu and Südhof, 2013). The specific role for RE in processing contextual, as opposed to discrete CSs, is consistent with previous work from our lab showing that HPC-PFC interactions are involved in contextual regulation of fear memories, rather than CS freezing *per se* (Jin and Maren, 2015a, 2015b; Wang et al., 2016; Marek et al., 2018). In contrast to the present results, Xu and Südhof (2013) did not observe deficits in the acquisition of contextual freezing after RE inactivation. However, as we have shown here, memories encoded under RE inactivation can be retrieved when the RE is offline. Irreversible inhibition of synaptic function in RE would allow hippocampal-independent memories of context to be encoded and expressed.

Unexpectedly, inactivating the RE prior to retrieval testing did not impair freezing in the conditioning context in animals that were conditioned intact. Previous work has shown that RE inactivation impairs memory retrieval in a variety of tasks (Hembrook and Mair, 2011; Hembrook et al., 2012; Cholvin et al., 2013; Jin and Maren, 2015a; Layfield et al., 2015), presumably by disrupting mPFC-HPC synchrony (Preston and Eichenbaum, 2013; Griffin, 2015; Jin and Maren, 2015a). However, as we have observed in the present study, RE inactivation does not always impair performance in hippocampal-dependent tasks (Loureiro et al., 2012). Indeed, a number of studies have observed spared retrieval of contextual fear memory after either muscimol (Holt and Maren, 1999; Maren and Holt, 2004; Matus-Amat et al., 2004; Sparks et al., 2011) or cobalt chloride (Resstel et al., 2008) infusions into the hippocampus, or optogenetic inhibition of dentate granule cells (Kheirbek et al., 2013; Bernier et al., 2017). Collectively, these data suggest that once contextual representations are learned (in an intact animal), they can be retrieved in the absence of the hippocampal system (Sparks et al., 2011).

The important role for RE in contextual fear conditioning is in line with previous work implicating the RE in HPC-dependent processes, such as spatial and working memory (Cassel and Pereira de Vasconcelos, 2015; Layfield et al., 2015; Hallock et al., 2016). In these tasks, memory relies heavily on contextual processing. During contextual fear conditioning, animals must encode a contextual representation that then comes into association with the aversive US. After acquisition, this memory yields conditioned freezing specific to the conditioning context and allows animals to discriminate the dangerous conditioning context from other, safe places. What might cause the contextual discrimination deficits observed after RE inactivation? One possibility is that animals without a functional RE use a non-hippocampal system to acquire an “elemental” representation of the context (Maren et al., 1997; Rudy and O’Reilly, 1999; Maren, 2001; Matus-Amat et al., 2004; Rudy and Matus-Amat, 2005). Bereft of a memory system that integrates multimodal sensory information into a unified, configural representation of context, animals with permanent damage to the RE might associate only the most salient features of the conditioning experience (perhaps even the experimenter or transport cues) with shock. These elemental associations would readily generalize across test contexts and fail to support subtle discriminations between contexts. Consistent with this view, many investigators have found that permanent hippocampal lesions fail to affect contextual conditioning *per se* (Maren et al., 1997; Frankland et al., 1998; Cho et al., 1999; Wiltgen et al., 2006), but impair contextual discrimination (Frankland et al., 1998; Cho et al., 1999).

Consistent with this proposition, we found that contextual memories acquired in animals after MUS infusions into the RE were not sensitive to intra-hippocampal infusions of APV. In contrast, contextual conditioning was impaired by APV infusions into DH in animals trained after SAL infusions into the RE, consistent with many other reports (Stiedl et al., 2000; Quinn et al., 2005; Matus-Amat et al., 2007; Czerniawski et al., 2012). These results indicate that RE inactivation forces animals to acquire contextual fear memories in a hippocampal-independent manner, presumably through alternate neural systems that associate contextual elements with the US (Maren et al., 1997; Rudy and O’Reilly, 1999; Barrientos et al., 2002; Matus-Amat et al., 2004; Rudy and Matus-Amat, 2005). It is clear that the amygdala mediates Pavlovian fear conditioning to both discrete and contextual CSs (Goosens and Maren, 2001; Maren, 2001; Wilensky et al., 2006), and it is likely that elements of the context (light, odor, or grid) are associated with the footshock US in the amygdala to support contextual conditioning in rats trained after RE inactivation. Previous studies have suggested that cortical regions such as prefrontal cortex, retrosplenial cortex, or perirhinal cortex can process contextual information independently of the hippocampus, and these regions may support context-US associations in the amygdala (Zelikowsky et al., 2013; Heroux et al., 2017; Coelho et al., 2018).

It has previously been argued that the hippocampal memory system, functions to acquire configural representations of context that enable context conditioning and discrimination (Maren et al., 1997; Frankland et al., 1998; Rudy and O’Reilly, 1999; Matus-Amat et al., 2004). Interestingly, in the absence of the hippocampal system, animals can acquire context memories using a non-hippocampal system that associates context elements with an aversive US to yield conditional freezing. However, this impoverished representation of context leads to considerable generalization, particularly across highly similar contexts. Under normal circumstances, hippocampal-dependent configural leaning overshadows the elemental learning system, which results in the encoding of context memories that later require the hippocampal system for retrieval (Sutherland et al., 2010; Sparks et al., 2011). In the context of the present results, this model suggests that the RE functions as a critical component of the hippocampal memory system involved in encoding precise contextual representations during fear conditioning. More specifically, our data reveal that 1) the RE functions to encode contextual memories that support contextual discriminations and 2) when conditioning occurs during RE inactivation, memories of conditioning are formed, but are only accessible when retrieved again under RE inactivation. Interestingly, we also show that RE inactivation does not prevent the retrieval of contextual memories *per se* and causes inappropriate and generalized fear to safe contexts. Altogether, these results are consistent with the proposal that conditioned freezing to an aversive context can be supported by either a configural representation encoded by the hippocampal system or by an elemental representation encoded outside the hippocampus (Maren et al., 1997; Rudy and O’Reilly, 1999; Anagnostaras et al., 2001; Chang and Liang, 2017; Sparks et al., 2011). Configural representations normally overshadow the elemental representation of context to dominate performance during retrieval. Similar to hippocampal lesions, RE inactivation impairs configural encoding of context representations, rendering an elemental memory of the conditioning experience that is inhibited when the hippocampal system is active at the time of retrieval.

By this account, animals conditioned after RE inactivation encode an elemental representation of context (e.g., context *A’*) that comes into association with the US and is only retrieved when the animal encounters context *A’* in the future. After fear conditioning, contextual freezing in animals conditioned under RE inactivation will only be expressed in context *A’*, which requires RE inactivation to be experienced. Moreover, this account assumes that configural representations of the context, which are encoded by the intact brain, are not only insufficient to retrieve the elemental *A’* association, but in fact inhibit its retrieval. Ultimately, this leads to both deficits in the contextual freezing and poor contextual discrimination. Overall, the present data reveal that the RE is critical for contextual processes involved in fear conditioning and suggest that it is a critical hub for HPC-mPFC interactions involved in learning and memory.

## Acknowledgements

This work was supported by grants from the McKnight Endowment for Neuroscience (McKnight Memory and Cognitive Disorders Award to SM) and the National Institutes of Health (R01MH065961 and R01MH117852) to SM. The authors would like to thank Kelsey J. Clement and Sohmee Kim for their technical assistance.

